# Pharmacological or genetic targeting of Transient Receptor Potential (TRP) channels can disrupt the planarian escape response

**DOI:** 10.1101/753244

**Authors:** Ziad Sabry, Alicia Ho, Danielle Ireland, Christina Rabeler, Olivier Cochet-Escartin, Eva-Maria S. Collins

## Abstract

In response to noxious stimuli, planarians cease their typical ciliary gliding and exhibit an oscillatory type of locomotion called scrunching. We have previously characterized the biomechanics of scrunching and shown that it is induced by specific stimuli, such as amputation, noxious heat, and extreme pH. Because these specific inducers are known to activate Transient Receptor Potential (TRP) channels in other systems, we hypothesized that TRP channels control scrunching. We found that chemicals known to activate TRPA1 (allyl isothiocyanate (AITC) and hydrogen peroxide) and TRPV (capsaicin and anandamide) in other systems induce scrunching in the planarian species *Dugesia japonica* and, except for anandamide, in *Schmidtea mediterranea*. To confirm that these responses were specific to either TRPA1 or TRPV, respectively, we tried to block scrunching using selective TRPA1 or TRPV antagonists and RNA interference (RNAi) mediated knockdown. Unexpectedly, co-treatment with a mammalian TRPA1 antagonist, HC-030031, enhanced AITC-induced scrunching by decreasing the latency time, suggesting an agonistic relationship in planarians. We further confirmed that TRPA1 in both species is necessary for AITC-induced scrunching using RNAi. Conversely, while co-treatment of a mammalian TRPV antagonist, SB-366791, also enhanced capsaicin-induced reactions in *D. japonica*, combined knockdown of two previously identified *D. japonica TRPV* genes (*DjTRPVa* and *DjTRPVb*) did not inhibit capsaicin-induced scrunching. Surprisingly, RNAi of either *DjTRPAa* or *DjTRPVa/DjTRPVb* disrupted scrunching induced by the endocannabinoid and TRPV agonist, anandamide. Overall, our results show that although scrunching induction can involve different initial pathways for sensing stimuli, this behavior’s signature dynamical features are independent of the inducer, implying that scrunching is a stereotypical planarian escape behavior in response to various noxious stimuli that converge on a single downstream pathway. Understanding which aspects of nociception are conserved or not across different organisms can provide insight into the underlying regulatory mechanisms to better understand pain sensation.

## Introduction

Normal locomotion of freshwater planarians, termed gliding, is achieved through synchronous beating of cilia in a layer of secreted mucus (1–3). Gliding planarians display a smooth motion without major body shape changes, except for turning movements of the anterior end. However, when exposed to certain noxious stimuli (e.g. low pH, high temperature, or amputation), planarians switch to a muscular-based escape gait that is characterized by oscillatory body length changes (4). We termed this gait scrunching and showed that it has a characteristic set of 4 quantifiable parameters: 1. frequency of body length oscillations, 2. relative speed, 3. maximum amplitude, and 4. asymmetry of body elongation and contraction (4). Moreover, scrunching is conserved among different planarian species, with each species demonstrating a characteristic frequency and speed. Although scrunching shares similarities with peristalsis, another muscle-based oscillatory gait that occurs when cilia beating is disrupted (2,5–7), scrunching is cilia-independent, can be induced in animals performing peristalsis, and is distinguishable from peristalsis based on the 4 parameters listed above (4), demonstrating that scrunching and peristalsis are distinct gaits. Because scrunching is such a stereotypical response involving many steps of neuronal communication (sensation, processing, neuro-muscular communication), scrunching in response to noxious heat has proven to be a useful and sensitive readout of neuronal function in planarian toxicological studies (8,9). However, which molecular targets and neuronal circuits regulate scrunching remain an open question.

Recently, it has been shown using RNA interference (RNAi) that the Transient Receptor Potential (TRP) channel, TRP ankyrin 1 (TRPA1), is required for avoidance behaviors in *Schmidtea mediterranea* in response to noxious heat and the pungent ingredient in mustard oil, allyl isothiocyanate (AITC) (10). The authors also showed that the noxious heat response was mediated by H_2_O_2_ and/or reactive oxygen species which directly activate TRPA1, causing the planarian to avoid hot regions. Moreover, in response to physical injury (tail snips), *Smed-TRPA1*-knockdown worms scrunched with a significantly reduced amplitude (10). Based on these results, the authors hypothesized that TRPA1 signaling, induced through H_2_O_2_ upregulation at the site of wounding, may regulate amputation-induced-scrunching in *S. mediterranea*. Whether H_2_O_2_ exposure alone induces scrunching or whether TRPA1 also plays a role in triggering scrunching in response to other stimuli is still unknown.

TRPA1 is a member of the TRP superfamily, comprised of widely conserved transmembrane, nonselective cation channels (11). TRP channels mediate responses to almost all classes of external stimuli, including various nociceptive stimuli such as extreme temperatures, UV light, and specific chemical irritants, and as such mediate the initial steps of pain sensation (11–13). TRPs are classified into sub-families depending on their main mode of activation (mechanical, thermal, chemical…), but are often polymodal, integrating different stimuli in the same channel (11,13,14).

In addition to TRPA1, TRPV channels are also good candidates for possibly regulating scrunching. Scrunching is activated by low pH and noxious heat (4), stimuli which are known to activate members of the TRP vanilloid (TRPV) sub-family, named after their sensitivity to vanilloid compounds such as capsaicin, in vertebrates and invertebrates (15–20). TRPV channels are activated by a diverse range of stimuli and exhibit a high level of species-dependent functional differences (20,21). For example, while human and rat TRPV1 are highly sensitive to capsaicin, rabbit and bird have greatly reduced sensitivities (21–23). While historically it was thought that, like fruit flies and nematodes (24–26), most invertebrates were also insensitive to this chili pepper irritant (27), medicinal leech was recently found to contain a capsaicin-sensitive TRPV channel (16). Interestingly, although no TRPV homologs exist in the parasitic flatworm *Schistosoma mansoni*, it was shown that TRPA1 in this species mediates the behavioral response to capsaicin (28,29). In the planarian *Dugesia japonica*, seven TRP genes have been previously cloned and their expression profiles characterized (30). TRPMa channels have been shown to regulate thermotaxis behavior at lower temperatures (0-25°C) (30), but the function of TRPA and TRPV channels has not yet been studied in this species.

Thus, based on previous literature in planarians and the known activators of scrunching, we hypothesized that TRPA1 and TRPV channels control planarian scrunching to specific stimuli. To test this hypothesis, we first assayed a variety of chemicals for their ability to induce scrunching in two freshwater planarian species, *S. mediterranea* and *D. japonica.* We focused on chemical compounds that have been shown to activate TRPA1 and/or TRPV, such as AITC, an agonist of planarian (10) and mammalian (31,32) TRPA1, and capsaicin, a specific agonist of mammalian TRPV1 (33,34).

We found that scrunching is a specific response to known modulators of TRPA1 or TRPV channels, including AITC and capsaicin, supporting the hypothesis that scrunching is controlled by TRPA1 and TRPV channels. These findings were substantiated by knocking down either *SmTRPA1, DjTRPAa* or *DjTRPVa/DjTRPVb* using RNAi and evaluating the behaviors of these worms when exposed to a subset of the confirmed scrunching inducers. We found that TRPA1 and TRPV elicit scrunching in response to different triggers and may have some overlapping functions.

The observation that scrunching is a stereotypical response that is the same for different stimuli and sensing mechanisms suggests the existence of a single convergent pathway that regulates scrunching downstream of TRP activation. TRP channels are involved in various chemical and physical sensing capacities across eukaryotes, from yeast to humans, yet exhibit a high level of diversity, both within the superfamily and across species (11,13). By understanding how these channels are used in different species, such as planarians, we gain better insight into their regulatory mechanisms, with the potential to reveal elements important to control pain sensation.

## Materials and methods

### Animal care

Asexual freshwater planarians of the species *Dugesia japonica* and *Schmidtea mediterranea* were used for all experiments. The animals were fed organic chicken or beef liver 1-2 times per week, cleaned twice per week, and starved for 5-7 days prior to experimentation. Planarians were stored in a temperature-controlled Panasonic incubator in the dark at 20°C with *D. japonica* in dilute (0.5 g/L) Instant Ocean (IO) water (Spectrum Brands, Blacksburg, VA, USA) and *S. mediterranea* in 1X Montjüic Salts (MS) water (35).

### Behavioral assays

#### Pharmacological perturbations

All chemicals used are listed in Table 1. Chemicals were stored according to supplier specifications. All stock solutions were made directly in IO or MS water or in 100% dimethyl sulfoxide (DMSO, Sigma-Aldrich, St. Louis, MO, USA), depending on chemical solubility. For chemicals prepared in DMSO, the final DMSO concentrations were kept ≤1%, which does not induce scrunching (S1 Fig). Specific conditions for each chemical experiment were determined empirically by qualitative observation of worm scrunching behavior (Table 1). The lowest exposure concentration tested which induced the most straight-line scrunches in wildtype worms was used for each experiment. The pH of all exposure solutions (except for hydrogen chloride) was checked and adjusted with NaOH to fall between 6.90-7.10, to ensure the observed scrunching behavior was not due to low pH conditions. Planarians were exposed to the chemicals either in the bathing solution or by pipetting a fixed volume directly onto a worm. Pipetting allowed for the usage of small volumes of locally higher chemical concentrations and was used in cases of poor chemical solubility or when baths failed to produce sufficiently long stretches of straight-line scrunching that could be used for quantitative analysis. Working solutions for chemicals were made fresh prior to starting experiments. Planarians were individually placed into 100 x 15-mm or 60 x 15-mm petri dishes (Celltreat Scientific Products, Shirley, MA, USA) depending on whether experiments were conducted in baths or by pipetting, respectively. The behavior of each planarian was recorded, starting immediately after initial exposure to the chemical, for up to five minutes at 10 frames per second (fps) using a charge-coupled device camera (PointGrey Flea3 1.3MP Mono USB 3.0) with a 16-mm lens (Tamron M118FM16 Megapixel Fixed-focal Industrial Lens) attached to a ring stand.

**Table 1.** Overview of chemicals used to induce scrunching.

| Chemical | CAS | Provider | Purity | Exposure concentration | Exposure method | Type/action and references |
| --- | --- | --- | --- | --- | --- | --- |
| Allyl isothiocyanate (AITC) | 57-06-7 | Sigma-Aldrich | 95% | Dj, Sm — 50, 75, 100 μM | 25 mL bath | TRPA1 agonist (10,31) |
| Hydrogen peroxide (H<sub>2</sub>O<sub>2</sub>) | 7722-84-1 | Sigma-Aldrich | 30% | Dj, Sm — 40 mM | 25 mL bath | TRPA1 agonist (10) |
| Capsaicin | 404-86-4 | Cayman Chemicals | ≥95% | Dj, Sm — 33, 82.5, 165 μM | 25 mL bath | TRPV1 agonist (15,20,31,33,34) |
| Anandamide | 94421-68-8 | Sigma-Aldrich | ≥97% | Dj – 100, 125 μM<br>Sm – 100 μM | 25 mL bath | Endocannabinoid and TRPV1 agonist (34,36) |
| HC-030031 | 349085-38-7 | Cayman Chemicals | ≥98% | Dj, Sm – 100 μM | 25 mL bath | TRPA1 antagonist (37–39) |
| SB-366791 | 472981-92-3 | Cayman Chemicals | ≥98% | Dj, Sm – 1, 10 μM | 25 mL bath | TRPV1 antagonist (16,40) |
| Hydrogen chloride (HCl) | 7647-01-0 | Sigma-Aldrich | 36.5-38.0% | Dj, Sm — pH to 2.7 | 10 μL pipette | Low pH scrunching inducer (4) |

#### Amputation experiments

Individual *S. mediterranea* planarians were placed into 100 x 15-mm petri dishes containing 15 mL of MS water. Using a razor blade, worms were amputated just above the pharynx. The behavior of each worm was recorded at 10 fps using the same setup as in the chemical assays. The number of scrunches of the resulting head piece was counted for each amputation with the scrunching sequence beginning after the first immediate contraction.

#### High temperature experiments

60mm x 15mm petri dishes were filled with 5 mL of either IO or MS water. Individual worms were placed in the dishes and induced to scrunch by pipetting 100 μL 65°C IO/MS as in (4). Additionally, we tested the effects of a heated water bath using an automated set-up for screening individual planarians in a 48-well plate (8). To induce scrunching, the plate was placed on a warmed peltier plate (TE Technology Inc., Traverse City, MI), whose temperature was computer controlled to heat the water in the wells. The temperature of the peltier was initially set to 65°C for the first 30 seconds to quickly heat up the plate from room temperature and then gradually decreased to 43°C to stabilize the aquatic temperature across the plate at around 32°C for 4 minutes. The plate was imaged from above and the movies were analyzed using a custom, automated MATLAB (MathWorks, Natick, MA, USA) script to detect instances of scrunching, as previously described (8).

#### Scrunching quantification

Recordings of planarian behavioral responses to the noxious stimuli were processed using ImageJ (National Institutes of Health, Bethesda, MD, USA). The background-subtracted image sequences were cropped to capture the first set of at least four consecutive straight-line scrunches or oscillations. An ellipse was then fit to the sequence of binary images of the worm to track and quantify the major axis (length of the worm) over time. From these data, the parameters frequency (number of scrunching/oscillation cycles per second), maximum elongation (difference between longest and shortest elongations/contractions as a fraction of the longest), relative speed (product of maximum elongation and frequency), and fraction of time spent elongating were then quantified in MATLAB as in (4). Unless stated otherwise, all values denote mean +/- standard deviation. Statistical significance for each scrunching parameter (or number of scrunches for amputation experiments) was calculated using a student’s t-test comparing to either previously published values for amputation for wild-type animals or to the *control* RNAi population for RNAi experiments.

#### Behavior scoring

Recordings of worms in chemical baths were scored by 2 blind reviewers. For every 15 s interval in the first 90 s of recording, worms were scored as either scrunching, exhibiting a non-scrunching reaction, or not reacting. Worms were scored as scrunching if they scrunched at least once in a given 15 s interval, based on the definitions set in (4). Worms were scored as exhibiting a non-scrunching behavior if they performed other behaviors, such as head shaking, frequent turning or abnormally long body elongation. Worms that glided unhindered throughout the 15 s interval were scored to have no reaction. Three experimental replicates were carried out for each condition, with N=8 *S. mediterranea* and N=10 *D. japonica* used per replicate. The mean of the scored responses from the two reviewers and across the experimental replicates for each 15 s interval are shown in the respective figures.

### Mucus staining

Staining procedures were performed using fluorescently labeled *Vicia villosa* (VVA) lectins as described previously (4). *D. japonica* planarians were individually placed in wells and induced to scrunch atop a glass coverslip using baths of 33 μM capsaicin or 50 μM AITC. The same staining procedure was followed for *S. mediterranea* planarians with scrunching being induced by a bath of 33 μM capsaicin or 50 μM AITC. Mucus trails were imaged in 4x under GFP fluorescence using a Nikon Eclipse Ci microscope (Nikon Corporation, Minato, Tokyo, Japan). Images were stitched together using Fiji (41) and the MosaicJ plugin (42).

### Cilia imaging

To view cilia beating, *D. japonica* and *S. mediterranea* planarians were incubated for five minutes in baths of 100 µM anandamide before mounting between a glass slide and a 22*22’ coverslip. Imaging procedures were performed as previously described in (4).

### Cloning of *Dugesia japonica* TRP genes

Partial mRNA sequences for TRPA1 (*DjTRPAa)* and TRPV (*DjTRPVa* and *DjTRPVb)* homologs in *D. japonica* have been previously published (30). Primers were designed using Primer3 (43) from these templates for *DjTRPVa* and *DjTRPVb* to generate 213 and 430 bp fragments, respectively (Table 2). For *DjTRPAa*, using the published sequence as a starting point, we blasted against a *D. japonica* transcriptome (dd_Djap_v4) on PlanMine (44) to identify the full coding sequence (transcript dd_Djap_v4_9060_1_1). An 895 bp fragment was identified from this transcript and cloned using the primers in Table 2.

**Table 2.**
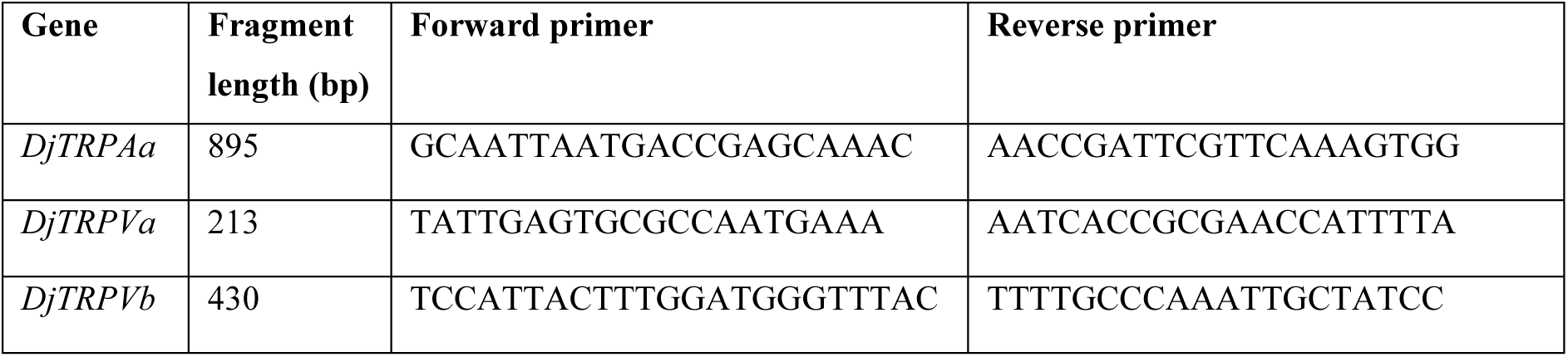
Primers used to clone *D. japonica* TRP genes.

These fragments were cloned into the pPR244-TRP vector using ligase independent cloning (45). The *Smed-TRPA1-pGEMt* plasmid was a gift from Dr. Marco Gallio (10).

### RNAi feedings and injections

Expression of *SmTRPA1 and DjTRPAa* were knocked down separately in their respective species. Expression of *DjTRPVa* and *DjTRPVb* were knocked down in combination and referred to as *DjTRPVab* RNAi. The respective TRP genes of interest were knocked down by injecting *S. mediterranea* or *D. japonica* worms on four consecutive days with *in vitro* transcribed dsRNA to a final concentration of at least 1 μg/μL as in (46). Negative control populations of both species, denoted as *control* RNAi, were injected with *unc22* dsRNA, a non-homologous *C. elegans* gene. Approximately 180 nL dsRNA were injected into each worm per day of injection using a standard dissection microscope and Pneumatic PicoPump Model PV 820 (World Precision Instruments, Sarasota, FL, USA). Needles were made by pulling 1-mm/0.58-mm OD/ID Kwik-Fill borosilicate glass capillaries through a two-stage program on a P-1000 micropipette puller (Sutter Instrument Company, Novato, CA, USA). On the fourth day of injection, after the fourth injection had been administered, worms were fed organic chicken liver mixed with at least 1 μg/μL dsRNA. Worms were then starved for six days prior to experiments.

### qRT-PCR

RNA was extracted from ten worms for each RNAi population using TRIzol (Invitrogen, Carlsbad, CA, USA) then purified using an RNeasy Mini Kit (QIAGEN, Germantown, MD, USA) including a DNase treatment. cDNA was synthesized from each RNA pool using the SuperScript® III First-Strand Synthesis System for RT-PCR (Invitrogen, Carlsbad, CA, USA), following the manufacturer’s protocol and priming with random hexamers. Primers for qPCR were designed using Primer3 (43) and are listed in Table 3.

**Table 3.**
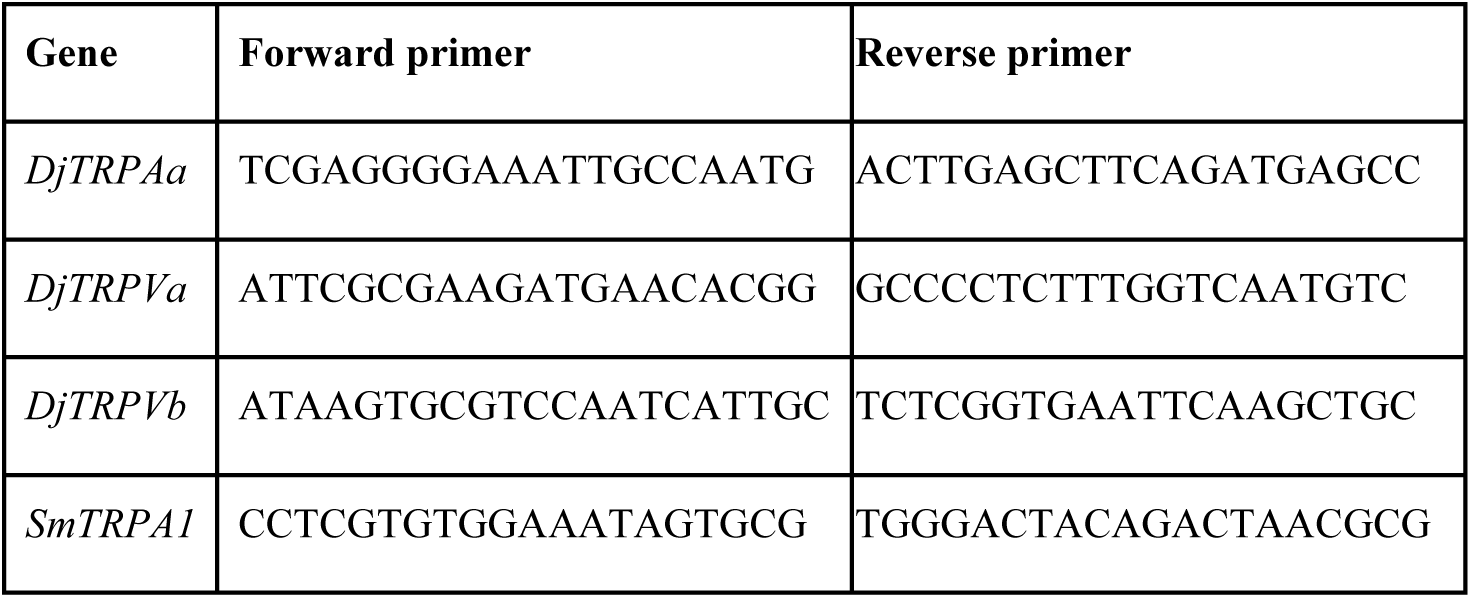
Primers used for qRT-PCR.

*DjGAPDH* and *SmedGAPDH* were used as housekeeping genes for their respective species. qPCR was performed on an MJ Research PTC-200 thermocycler equipped with a Chromo4 Real-Time PCR Detector (Bio-Rad Laboratories, Hercules, CA, USA), using PerfeCTa® SYBR® Green FastMix® (Quantabio, Beverly, MA, USA). Technical triplicates were run for all reactions within an experiment, and two biological replicates were performed. To analyze primer efficiency, standard curves were obtained using a 1:1:1:1 mix of all cDNA pools for each species, serially diluted. The efficiency for each primer pair was found to be between 87 - 116%. Analysis of relative expression for the genes targeted by RNAi was performed using the ΔΔCt method, where reported values are the mean of all replicates.

## Results and discussion

### TRPA1 and TRPV agonists induce scrunching in *S. mediterranea* and *D. japonica*

Based on the known inducers of scrunching in *D. japonica* and *S. mediterranea* (noxious heat, pH, amputation) ((4) and S2 Fig), and the recent work by Arenas et al. suggesting that TRPA1 mediates scrunching in response to amputation in *S. mediterranea* (10), we tested known chemical agonists of planarian and other species’ TRPA1 and TRPV (Table 1) for their ability to trigger scrunching in *D. japonica* and *S. mediterranea*. The two oscillatory planarian gaits, scrunching and peristalsis, can be hard to distinguish qualitatively by eye. Thus, if oscillatory motion was observed, we distinguished peristalsis and scrunching by quantifying the 4 characteristic parameters (frequency, speed, maximum elongation, asymmetry of elongation/contraction) and comparing with published reference values for these gaits (4). We previously found that each planarian species exhibits a characteristic scrunching frequency and speed, with *D. japonica* scrunching at higher speeds and with almost double the frequency of *S. mediterranea* planarians (4). Therefore, all comparisons are done with references in the same species.

Under normal conditions, planarians glide, maintaining a constant body length over time (Fig 1A, B). When exposed to 50 µM of the TRPA1 activator, AITC, planarians scrunched showing oscillations of body length elongation and contraction (Fig 1C) with quantitative parameters consistent with those previously determined for *D. japonica* and *S. mediterranea* using amputation (Table 4).

**Fig 1.**
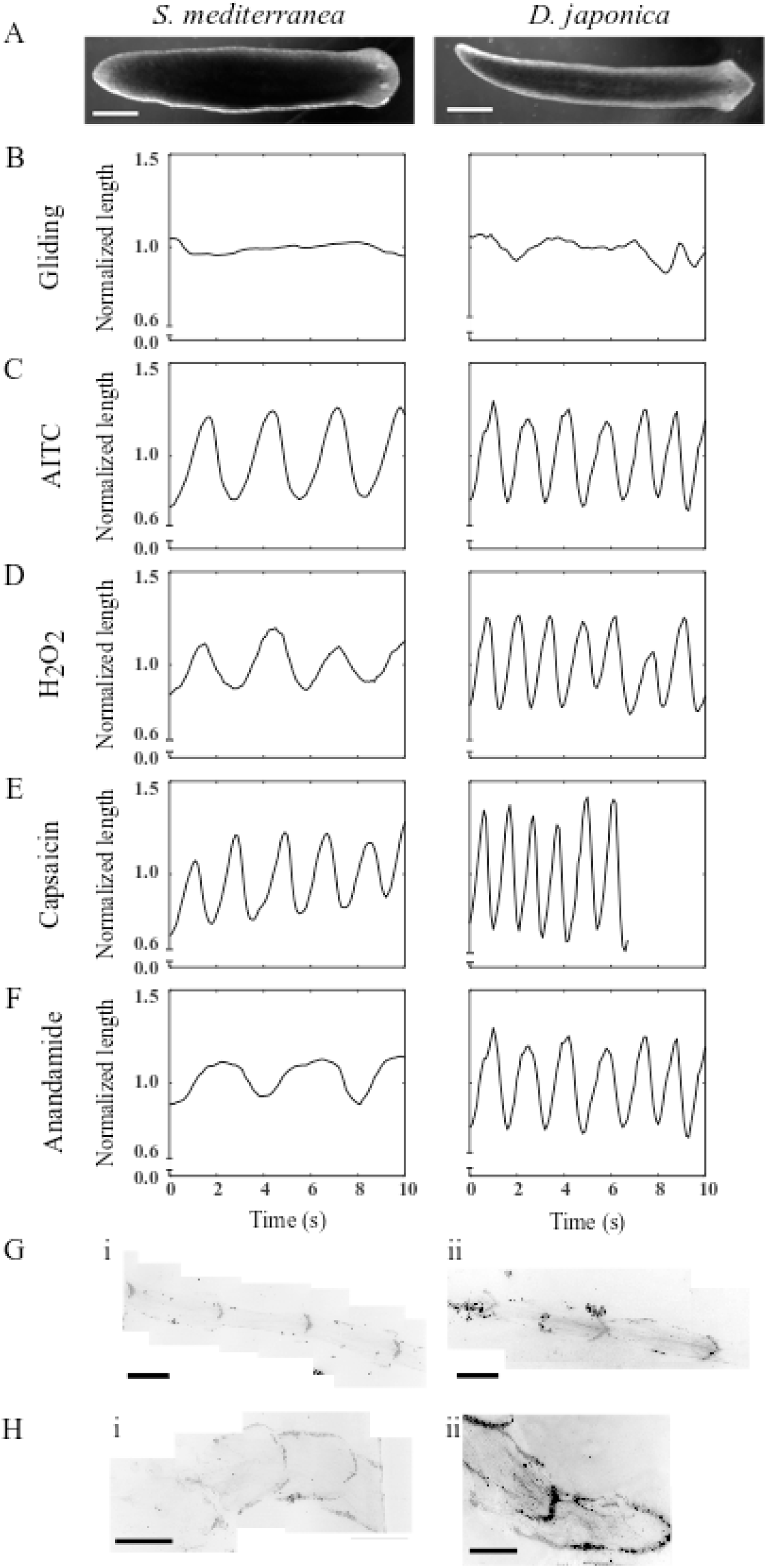
TRPA1 and TRPV agonists induce scrunching in planarians. (A) Single frames of gliding *S. mediterranea* (left) and *D. japonica* (right). (B-F) Representative length versus time plots for *S. mediterranea* (left) and *D. japonica* (right) planarians during (B) gliding or (C-F) chemically induced scrunching, when exposed to (C) 50 μM AITC, (D) 40 mM H_2_O_2_, (E) 165 μM capsaicin, or (F) 100 μM anandamide. Length was normalized by the average gliding length for all plots. (G-H) Example *S. mediterranea* (i) and *D. japonica* (ii) mucus trails stained with fluorescein-conjugated VVA lectin (see Methods) for worms exposed to (G) 50 μM AITC or (H) 165 μM capsaicin. Mucus trail images were black/white inverted. Scale bars: 1 mm.

**Table 4.** Quantification of scrunching parameters in response to TRPA1 and TRPV chemical agonists in *D. japonica* and *S. mediterranea*.

| Species | Stimulus | Conc. | Frequency (cycles s<sup>-1</sup>) | Maximum elongation | Speed (body length s<sup>-1</sup>) | Fraction of time spent elongating | N |
| --- | --- | --- | --- | --- | --- | --- | --- |
| *D. japonica* | AITC | 50 μM | 0.72 ± 0.08 | 0.52 ± 0.04 | 0.37 ± 0.04 | 0.59 ± 0.03 | 9 |
| *D. japonica* | H<sub>2</sub>O<sub>2</sub> | 40 mM | 0.54 ± 0.09 | 0.44 ± 0.04 | 0.23 ± 0.05<sup>^</sup> | 0.60 ± 0.02 | 8 |
| *D. japonica* | Capsaicin | 165 μM | 0.80 ± 0.14 | 0.48 ± 0.07 | 0.38 ± 0.05 | 0.59 ± 0.04 | 8 |
| *D. japonica* | Anandamide | 100 μM | 0.63 ± 0.08 | 0.43 ± 0.05<sup>^</sup> | 0.27 ± 0.05 | 0.57 ± 0.03 | 8 |
| *D. japonica* | Amputation<sup>a</sup> | -- | 0.70 ± 0.27 | 0.50 ± 0.08 | 0.34 ± 0.12 | 0.60 ± 0.12 | 15 |
| *S. mediterranea* | AITC | 50 μM | 0.33 ± 0.02 | 0.43 ± 0.05 | 0.14 ± 0.02 | 0.58 ± 0.02 | 5 |
| *S. mediterranea* | H<sub>2</sub>O<sub>2</sub> | 40 mM | 0.37 ± 0.03 | 0.39 ± 0.05 | 0.14 ± 0.02 | 0.52 ± 0.02 | 10 |
| *S. mediterranea* | Capsaicin | 165 μM | 0.44 ± 0.04 | 0.49 ± 0.06 | 0.21 ± 0.03 | 0.57 ± 0.03 | 8 |
| *S. mediterranea* | Anandamide | 100 μM | 0.28 ± 0.03<sup>^^</sup> | 0.30 ± 0.05<sup>^^</sup> | 0.08 ± 0.01<sup>^^</sup> | 0.49 ± 0.03<sup>^^</sup> | 7 |
| *S. mediterranea* | Amputation<sup>a</sup> | -- | 0.40 ± 0.09 | 0.44 ± 0.09 | 0.17 ± 0.09 | 0.62 ± 0.18 | 77 |
| *S. mediterranea* | Peristalsis<sup>a</sup> | -- | 0.26 ± 0.07 | 0.23 ± 0.19 | 0.06 ± 0.04 | 0.50 ± 0.07 | 14 |

Values denote mean ± std. All statistical analyses were conducted against the published amputation data.
<sup>^</sup> p < 0.05 and <sup>^^</sup> p < 0.01.
<sup>a</sup>Amputation and peristalsis data are previously published values (4), provided for reference.

Previous studies in *S. mediterranea* demonstrated that TRPA1 is directly activated by H_2_O_2_ (10); therefore, we assayed whether H_2_O_2_ exposure could induce scrunching. As expected from the results of AITC exposure, 40 mM H_2_O_2_ elicited scrunching in both planarian species (Fig 1D and Table 4). Notably, while *S. mediterranea* scrunching parameters were statistically insignificant from those induced by amputation, *D. japonica* scrunched at slightly (but statistically significant) lower speeds in 40 mM H_2_O_2_ compared to the reference values for amputation-induced scrunching (Table 4). This decrease in speed may be due to negative health effects of the exposure, since we found that *D. japonica* but not *S. mediterranea* disintegrated within a day following the 5 min H_2_O_2_ exposure unless excessively washed post H_2_O_2_ exposure. Even after washing in three separate 50 mL baths of IO water, 1/12 *D. japonica* planarians died within 24 hours. Additional range-finding tests were unable to determine a concentration of H_2_O_2_ that induced scrunching without negative health effects in *D. japonica.* Lower concentrations (10 and 20 mM) only induced head wiggling within 5 min of exposure. It is currently unclear why *D.* japonica show such an increased sensitivity to H_2_O_2_.

We have previously demonstrated that scrunching is induced by high heat and low pH (4), which are known activators of TRPV1 in other species (15–20). Thus, we tested whether the classical TRPV1 agonist, capsaicin, and the endocannabinoid anandamide, known to directly activate TRPV1 in other systems (34,36,47,48), were able to induce scrunching (Fig 1E, F). Both *S. mediterranea* and *D. japonica* scrunched with stereotypical parameters when exposed to 165 µM capsaicin (Fig 1E and Table 4). Additionally, in both species, scrunching in capsaicin was often accompanied by vigorous head shaking and jerking (S1 Movie).

In contrast, exposure to 100 μM anandamide elicited scrunching in *D. japonica* but not in *S. mediterranea* (Fig 1F and Table 4). *S. mediterranea* worms displayed oscillatory locomotion (Fig 1F), but a quantitative analysis of the parameters shows that *S. mediterranea* performed peristalsis (Table 4), which we have previously demonstrated to be a distinct gait from scrunching (4). Peristalsis is induced when cilia beating is disrupted, whereas scrunching is cilia-independent (2,4–7). Using cilia imaging, we confirmed that cilia beating was disrupted in *S. mediterranea* but not *D. japonica* exposed to 100 μM anandamide (S3 Fig). A possible explanation for this finding is that in addition to being a low potency agonist of TRPV1, anandamide also activates cannabinoid receptor 1 (CB-1). Although the cannabinoid receptor(s) have not yet been directly identified in planarians, pharmacological experiments with specific cannabinoid receptor agonists and antagonists in the planarian *Dugesia gonocephala* suggest the presence of functional cannabinoid receptor homologs in planarians (49,50). Complicating matters, in other systems, crosstalk with the cannabinoid system can modulate the responsiveness of TRPV1 (36). Furthermore, the efficiency of anandamide binding to TRPV1 appears to be species-specific (34). Thus, these factors could interact to produce different manifestations of similar, yet distinct, oscillatory gaits in the two species, resulting in scrunching in *D. japonica,* but peristalsis in *S*. *mediterranea*. This dissimilarity in behavioral phenotypes, together with the sensitivity differences to H_2_O_2_ exposure, emphasize that care needs to be taken when attempting to extrapolate findings from pharmacological studies from one planarian species to another.

Finally, we also visualized the mucus trails of worms of both species exposed to either AITC or capsaicin (Fig 1G, H) and saw the characteristic profiles of triangular anchor points that we have previously demonstrated to be associated with scrunching (4).

In summary, we found that archetypal agonists of TRPA1 and TRPV channels induce scrunching in planarians, supporting our hypothesis that induction of this gait is mediated by TRPA1 and TRPV activation.

### Increasing concentrations of AITC and the TRPA1 antagonist HC-030031 enhance scrunching

It has been previously shown that increasing the concentration of AITC decreases the latency to initiate the nocifensive response in the medicinal leech (39). We observed this same trend in both planarian species, as at higher concentrations of AITC more planarians scrunched earlier (Fig 2A), with more pronounced differences seen in *S. mediterranea*. However, in all concentrations of AITC tested, even when planarians did not scrunch initially, they still reacted to the AITC as evidenced by vigorous head turning (shown as percent worms reacting in Fig 2A), thus demonstrating that the initial sensation of AITC does not appear to be affected by concentration within the range tested here. Interestingly, the scrunching parameters were also dependent on AITC concentration, with increasing concentrations causing significantly increased maximum elongation and speed for both *D. japonica* and *S. mediterranea* when compared to 50 µM AITC (S1 Table). Whereas for *D. japonica*, these values, as well as frequency in 100 µM AITC, were also significantly different from the scrunching parameters in response to amputation, the parameters for *S. mediterranea* exposed to the different concentrations of AITC were not significantly different from amputation-induced scrunching parameters, indicating these differences were within the range observed in this species (S1 Table).

**Fig 2.**
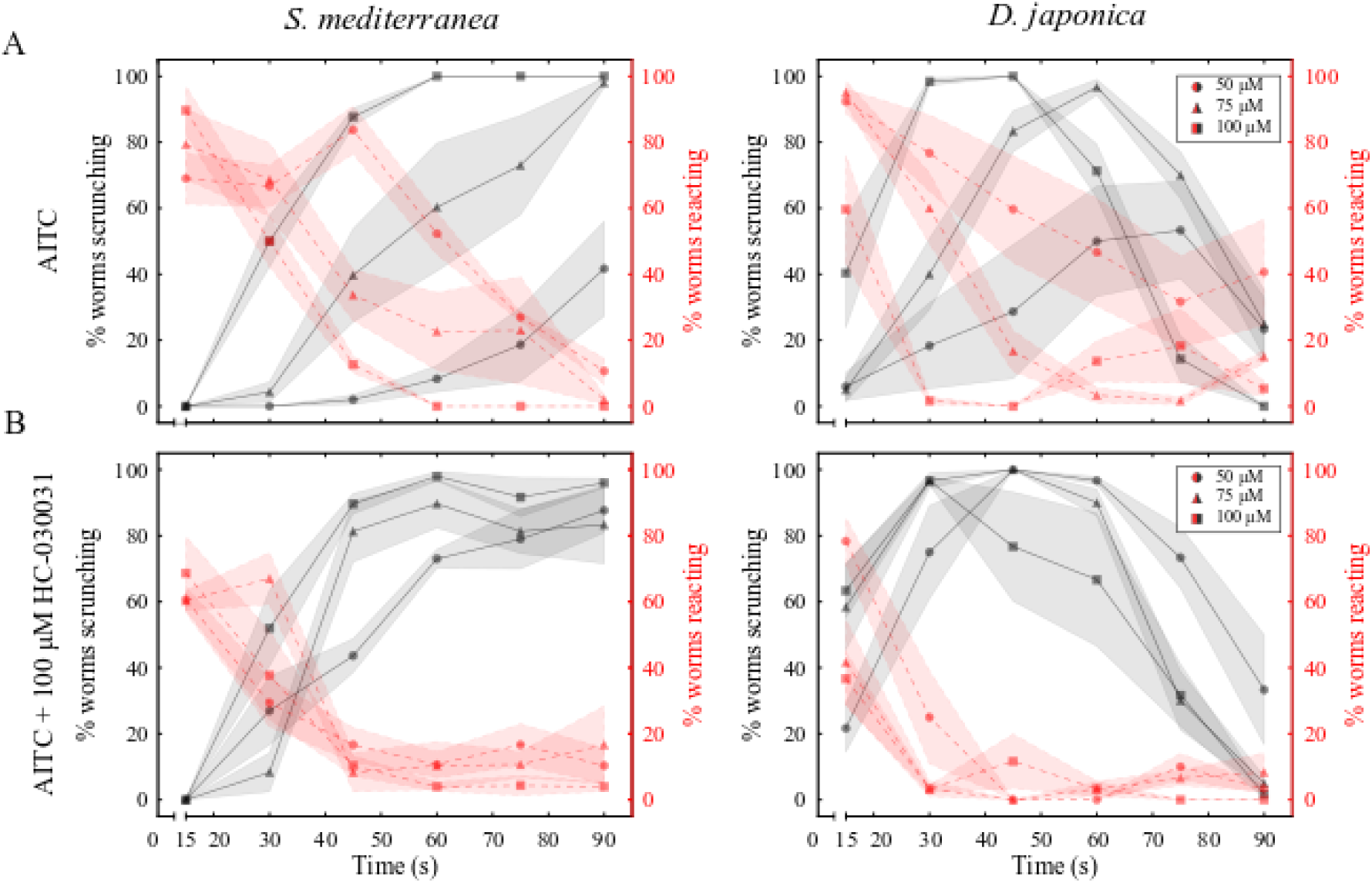
HC-030031 decreases scrunching latency induced by AITC. (A, B) Behavior scoring plots for *S. mediterranea* (left) and *D. japonica* (right) showing the percentage of worms scrunching (black lines) or reacting (behaviors other than scrunching, see Methods; red lines) every 15 s over 90 s when exposed to (A) AITC or (B) AITC + 100 μM HC-030031. Markers and shading represent the mean and standard deviation of 3 technical replicates, respectively.

A striking behavioral difference was observed between *S. mediterranea* and *D. japonica* when exposed to AITC. In all tested AITC concentrations, the majority (at least ∼80% in all tested concentrations) of scrunching *D. japonica* ceased scrunching by 90 seconds and began gliding, as seen by the decrease in both the percent worms scrunching and reacting in Fig 2A. This apparent desensitization was concentration-dependent; *D. japonica* planarians at higher AITC concentrations started and ceased scrunching earlier than those at lower AITC concentrations. *S. mediterranea* did not share this behavior and showed longer periods where all worms were scrunching (Fig 2A, compare 100 µM AITC between the two species). Consistent with this observed desensitization to prolonged scrunching activation in *D. japonica*, the continuous application of high concentrations of AITC completely desensitizes currents in the dorsal root ganglion neurons of mice (51).

HC-030031 is a specific TRPA1 antagonist that has been shown to block nocifensive responses to AITC in other systems, including rat and the medicinal leech (37,39). Therefore, we tested whether HC-030031 could block or at least attenuate planarian scrunching. During initial tests with multiple concentrations of HC-030031, we unexpectedly found that 200 μM HC-030031 induced scrunching in 10/10 *D. japonica* planarians at some point within 2 minutes of exposure, whereas it did not have that effect on *S. mediterranea* (S4A Fig). At 100 μM HC-030031, neither planarian species scrunched, but *D. japonica* displayed a mild reaction including vigorous head turning, which was not observed in *S. mediterranea*. However, since scrunching was absent at this concentration in both species, 100 μM HC-030031 was used for further experiments.

When co-administered with 50, 75, or 100 μM AITC, 100 μM HC-030031 decreased the latency to induce scrunching in the majority of planarians at 50 and 75 µM AITC in both planarian species (Fig 2, S2-3 Movies) suggesting a cooperative interaction between AITC and HC-030031, which mimicked the trend seen in increasing concentrations of AITC alone (Fig 2A). This effect was not as pronounced at 100 µM, suggesting that the maximal activity may have already been reached with 100 μM AITC.

Together, these findings suggest that increasing concentrations of AITC or cooperative actions of AITC and HC-030031 enhance scrunching, further supporting the idea that TRPA1 is involved in mediating the scrunching response.

### Genetic modulation of TRPA1 expression disrupts scrunching in response to AITC, H_2_O_2_, and amputation

Our chemical experiments suggest that TRPA1 activation in *S. mediterranea* and *D. japonica* induces scrunching. To confirm this, we knocked down *SmTRPA1* and *DjTRPAa* using RNAi and evaluated how this affected scrunching in response to TRPA1 modulators. Gene knockdown was confirmed by qRT-PCR showing 73.0% knockdown in *SmTRPA1* RNAi populations and 51.4% knockdown in *DjTRPAa* RNAi populations compared to expression in the species-specific *control* RNAi populations (S5A-B Fig).

When exposed to 100 μM AITC for 90 s, none of the *SmTRPA1* RNAi or *DjTRPAa* RNAi planarians scrunched, while all *control* RNAi animals of each species scrunched under the same conditions (Fig 3A-B, S4-5 Movies). Similarly, scrunching in response to 200 μM HC-030031 alone or to 100 μM HC-030031 + 100 μM AITC was completely lost in *DjTRPAa* RNAi animals (S4B-C Fig), demonstrating that TRPA1 is essential for AITC-induced scrunching and that HC-030031 activates TRPA1. Our results suggest a cooperative, rather than antagonistic, action of HC-030031 on planarian TRPA1. Different organisms have been shown to have different sensitivities to the antagonistic effects of HC-030031. For example, divergence of a single amino acid (N855 in human TRPA1) in frog and zebrafish TRPA1 is responsible for their insensitivity to the inhibitor (52). Although the mechanism of TRPA1 inhibition by HC-030031 could not be resolved structurally (53), it has been suggested to cause a conformational change in TRPA1 which disrupts ligand binding (52). Thus, it is possible that in planarians HC-030031 may cause a different conformational change in TRPA1 to instead potentiate AITC activation.

**Fig 3.**
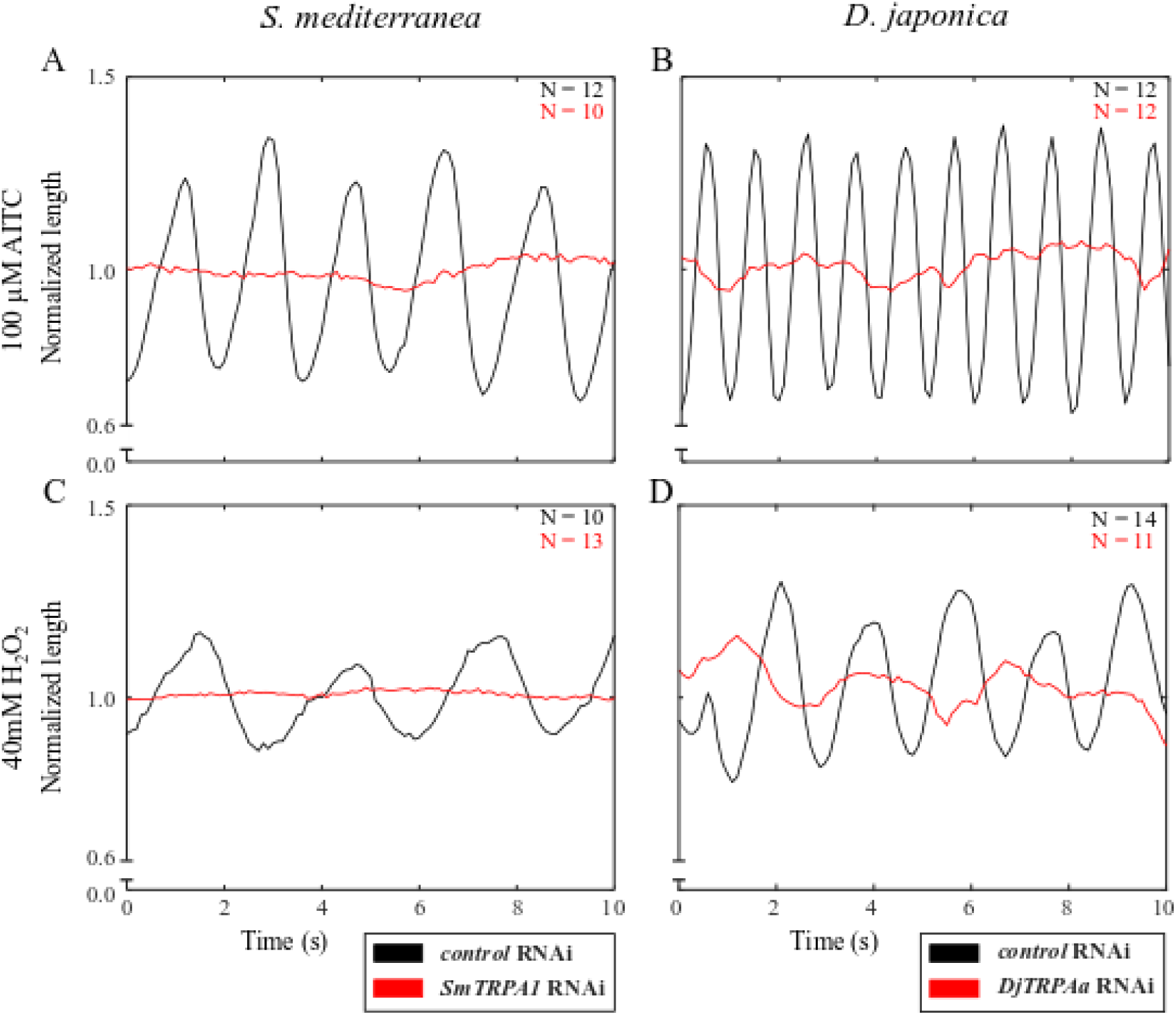
TRPA1 is necessary for AITC- and H_2_O_2_-induced scrunching. (A-D) Representative length versus time plots for RNAi treated *S. mediterranea* (left) and *D. japonica* (right) exposed to (A-B) 100 μM AITC or (C-D) 40 mM H_2_O_2_. Note that the scrunching frequencies in the *control* RNAi populations differ between the two inducers because a higher (100 μM) concentration of AITC was used (compare values in Table 4 and S1 Table). Plots are representative of the total number of worms tested, as indicated in the respective panels for each condition.

Scrunching was also completely lost in both *SmTRPA1* RNAi and *DjTRPAa* RNAi populations exposed to 40 mM H_2_O_2_ for either 270 s or 60 s, respectively, during which times all *control* RNAi planarians of both species scrunched (Fig 3C-D). These data are consistent with previous reports of H_2_O_2_ as a direct activator of *S. mediterranea* TRPA1 (10). Together, these results show that TRPA1 is essential to induce scrunching with either AITC or H_2_O_2_.

One of the most robust but unspecific inducers of scrunching is amputation (4). Arenas et al. found that when doing tail snips on filter paper, *Smed-TRPA1* RNAi animals exhibited a decreased scrunching amplitude compared to *control* RNAi animals (10). Because dry environments alone can induce scrunching (S6A Fig and (4)), we did not perform amputation experiments on filter paper, but in an aqueous environment instead. Under these experimental conditions, we found that knockdown of *SmTRPA1* caused reduced scrunching compared to *control* RNAi animals after amputation, evidenced by fewer total scrunches (S6B Fig). Thus, SmTRPA1 appears to partially mediate scrunching in response to amputation. These results are consistent with the work of Arenas et al., who also found attenuated rather than completely abolished scrunching in *Smed-TRPA1* RNAi animals after amputation (10).

In *S. mediterranea*, it has been shown that amputation leads to a burst of H_2_O_2_ production at the wound site (54). Thus, because H_2_O_2_ directly activates SmTRPA1, it has been suggested that mechanical injury (such as amputation) indirectly activates SmTRPA1 through H_2_O_2_ production (10). While our results confirm that H_2_O_2_ activation of TRPA1 induces scrunching, H_2_O_2_ activation of TRPA1 is likely not the only mechanism mediating amputation-induced scrunching since scrunching is not completely abolished in amputated *SmTRPA1* RNAi planarians. We were unable to perform these same experiments with the *D. japonica* RNAi populations as even in *control* RNAi animals, amputation only induces few scrunches robustly.

Together, our results confirm that TRPA1 in both *S. mediterranea* and *D. japonica* is necessary to induce scrunching in response to AITC and H_2_O_2_ and is partially involved in amputation-induced scrunching in *S. mediterranea*.

### TRPV antagonist SB-366791 enhances scrunching

As we did for TRPA1, we similarly dissected the role of TRPV in scrunching. Because anandamide did not elicit scrunching in *S. mediterranea* and because the *D. japonica* TRPV genes have previously been characterized (30), we carried out all further experiments in *D. japonica* only. When comparing the behavioral responses to different concentrations of capsaicin, we found that at all tested concentrations, the planarians initially reacted by vigorously turning and shaking their heads and then transitioned to a scrunching phenotype over time. Because of this transitional nature of the behavior, it was often difficult to confidently distinguish non-scrunching reactions from scrunching by eye alone. Increasing the concentration of capsaicin decreased the latency time to switch to a scrunching reaction, similarly to AITC (Fig 4A). As with AITC (Fig 2A), after 45 s in 165 μM capsaicin, *D. japonica* scrunching behavior began to cease (Fig 4A). However, unlike in AITC where *D. japonica* began gliding again, *D. japonica* continued to react in capsaicin by maintaining a contracted body length and exhibiting minor oscillations (S6 Movie). Many nociceptors, including TRPA1 and TRPV, have been shown to become desensitized following prolonged activation. This desensitization is why certain TRP agonists, such as capsaicin, have been used therapeutically as analgesics (55). In rat neuronal cell culture, it was shown that prolonged capsaicin exposure causes TRPV1 channels to be removed from the membrane through endocytosis and lysosomal degradation (56). A similar desensitization to scrunching induction appears to be present in *D. japonica*, though the underlying mechanism remains to be determined.

**Fig 4.**
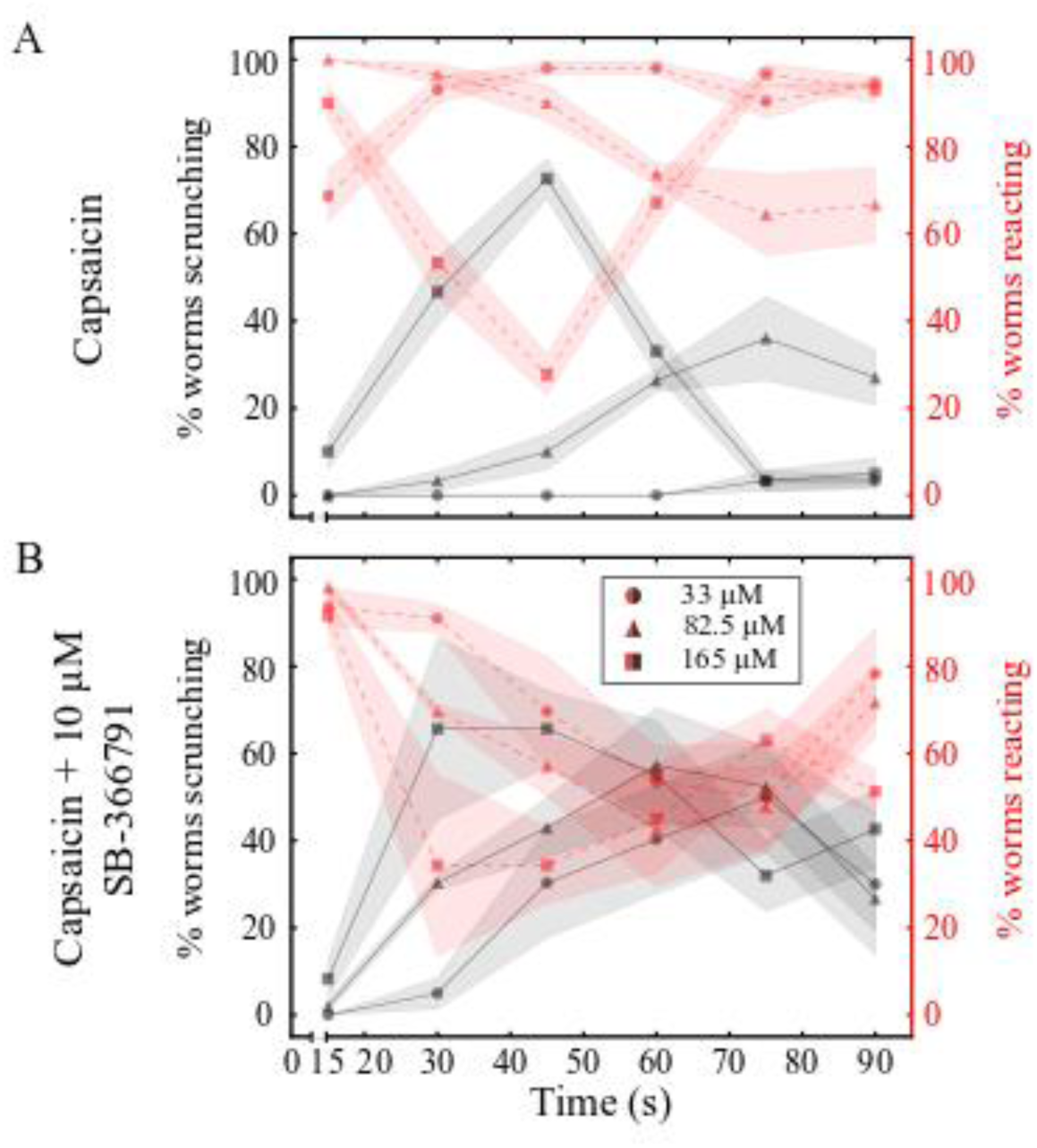
Scrunching in capsaicin is enhanced by SB-366791. (A-B) Behavior scoring plots for *D. japonica* showing the percentage of worms scrunching (black lines) or reacting (non-scrunching behaviors, red lines) every 15 s over 90 s when exposed to (A) capsaicin or (B) capsaicin + 10 μM SB-366791. Markers and shading represent the mean and standard deviation of 3 technical replicates, respectively.

We then tested a TRPV-selective antagonist SB-366791 (40). Initial experiments using a range of different concentrations of SB-366791 showed that *D. japonica* began vigorous head turning at a concentration of 10 μM SB-366791 but did not scrunch. No abnormal behaviors were observed at 1 μM (S7 Movie). Similarly to AITC co-administration with HC-030031, co-administration of capsaicin with 10 μM SB-366791 decreased the scrunching latency (Fig 4B). However, unlike the trends seen with HC-030031, co-administration with SB-366791 decreased the number of worms which stopped scrunching over time, creating a more prolonged scrunching reaction compared to capsaicin alone. Together these results suggest that although SB-366791 enhances capsaicin-induced scrunching it does not have the same potentiation effects seen with HC-030031 and AITC.

### *DjTRPVab* and *DjTRPAa* modulate scrunching behavior in response to anandamide

To assay their roles in mediating scrunching in response to the TRPV modulators, we knocked down both known *DjTRPV* genes (*DjTRPVa* and *DjTRPVb*) (30) in combination via RNAi (referred to as *DjTRPVab* RNAi). Gene knockdown was confirmed by qRT-PCR showing an 41.2% decrease in *DjTRPVa* and 83.3% decrease in *DjTRPVb* in the *DjTRPVab* RNAi population compared to expression in *control* RNAi planarians (S5C-D Fig). Because TRPA1 has been found to modulate sensitivity to capsaicin in the parasitic flatworm *S. mansoni*, which does not have any TRPV homologs (28,29,57), we also evaluated the reactions of *DjTRPAa* RNAi worms to the TRPV agonists. Neither *DjTRPVab* nor *DjTRPAa* RNAi ablated scrunching in response to 165 μM capsaicin (Fig 5A and S6 Movie). Although there was a slight decrease in the percentage of worms scrunching in the *DjTRPVab* and *DjTRPAa* RNAi populations compared to *control* RNAi, all *DjTRPVab* and *DjTRPAa* RNAi planarians either reacted or scrunched when exposed to 165 μM capsaicin. Similarly, for *S. mediterranea*, pipetting 165 μM capsaicin onto *control* or *SmTRPA1* RNAi populations induced scrunching in all tested animals (N=10). A TRPV homolog has not yet been identified in this species. Thus, none of these channels are solely responsible for capsaicin-induced scrunching.

**Fig 5.**
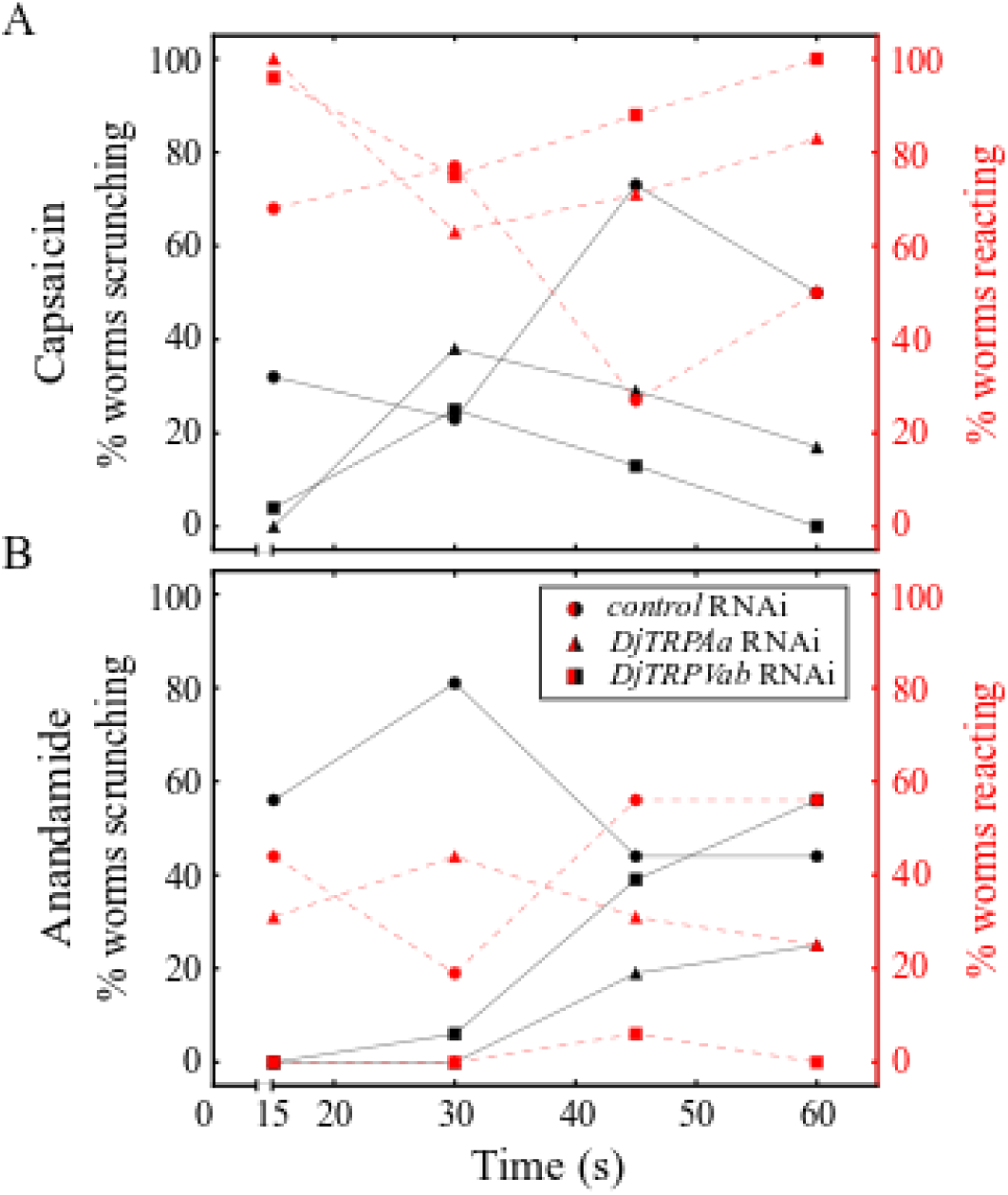
DjTRPAa and DjTRPV mediate scrunching in response to anandamide. (A-B) Behavior of *control* RNAi, *DjTRPAa* RNAi, and *DjTRPVab* RNAi planarians in (A) 165 μM capsaicin (N=11-12) and (B) 125 μM anandamide (N=8-9).

Although mammalian TRPV1 was originally identified as the “capsaicin receptor”, capsaicin-sensing ability and the responsible receptor varies dramatically across invertebrates. Several invertebrate species, including fruit flies and nematodes, are insensitive to capsaicin (25,26). In *Caenorhabditis elegans*, capsaicin potentiates the thermal avoidance response, but this effect is not dependent on OSM-9, the purported *C. elegans* TRPV1 homolog, suggesting another unknown receptor is involved (25). A similar situation appears to be present in *D. japonica*, where scrunching is not dependent on the identified TRPV homologs, DjTRPVa and DjTRPVb (Fig 5A). Our results also show that, unlike in *S. mansoni*, TRPA1 in both planarian species is not responsible for capsaicin sensing, suggesting evolutionary divergence. Together, these data suggest another receptor is likely responsible for mediating scrunching in response to capsaicin. What receptor is responsible for capsaicin sensing, whether it be another unidentified TRPV, a different TRP channel, or some unrelated protein, in freshwater planarians remains to be determined.

Next, we tested whether scrunching induced by anandamide could be affected by knockdown of either *DjTRPVab* or *DjTRPAa.* When exposed to 125 μM anandamide, most *control* RNAi planarians started scrunching within 15 s and continued scrunching for up to 60 s, with the remaining proportion exhibiting vigorous head turning (Fig 5B, S8 Movie). In both *DjTRPVab* or *DjTRPAa* RNAi planarians, anandamide-induced scrunching was attenuated, evidenced by an overall decrease in the percentage of worms scrunching and an increase in the latency time to induce scrunching (Fig 5B, S8 Movie). Thus, these data suggest some overlapping functions of DjTRPAa and DjTRPVab to mediate anandamide-sensing in *D. japonica.* Similar overlapping or interacting functions of TRPA1 and TRPV1 have previously been suggested in several other systems, including mouse (31), nematodes (58) and the medicinal leech (39).

Anandamide and other cannabinoids have complicated relationships with TRP channels. Both endogenous and synthetic cannabinoids act through the canonical cannabinoid receptors, CB-1 and CB-2, but some have been found to activate TRPV and TRPA1 channels as well (59,60). Particularly, in addition to its effects on TRPV1, anandamide has been shown to directly activate rat TRPA1(61), consistent with our findings that *DjTRPAa* RNAi planarians show attenuated scrunching in response to anandamide. Additionally, because of the extensive crosstalk between the endocannabinoid system and TRPV1, leading to sensitization of TRPV1 to other endogenous ligands, it has been suggested that even when anandamide treatment mimics the physiological outcomes of TRPV agonists, the effects are not necessarily due to direct activation of TRPV1 (36). Thus, it is unclear from our RNAi results whether anandamide’s role in scrunching is due to direct activation of TRPV (and/or TRPA1) or indirectly through its role as an endocannabinoid.

Finally, we assayed the scrunching response of all RNAi populations to noxious heat and low pH exposure, which are known to affect TRPV in other species (15,16,18–20). We observed scrunching in all populations (S7 Fig), indicating that none of these 3 genes are involved in the scrunching response to these stimuli. Strengthening this conclusion is the observation that, when scrunching was induced by heating the aquatic environment, scrunching was still observed in all RNAi populations with no statistically significant differences determined by a Fisher’s exact test with a p-value of 0.05. Scrunching was found in 24/34 *DjTRPAa* and 20/28 *DjTRPVab* RNAi planarians, similar to *control* RNAi worms (25/36). Consistent results were also found for *S. mediterranea* as similar proportions of animals scrunched in the *control* (9/22) or *SmTRPA1* (7/21) RNAi populations. The finding that in this assay *S. mediterranea* planarians across RNAi populations scrunched much less than *D. japonica* planarians may be a consequence of the experimental setup being optimized for *D. japonica* (8). While previous reports have shown that SmTRPA1 is involved in mediating heat avoidance behaviors via direct activation by H_2_O_2_ (10), our data suggest that other channels may be involved in triggering scrunching in response to high temperatures.

Taken together, these data demonstrate that TRPA1 mediates scrunching in response to AITC and H_2_O_2_ in both planarian species, whereas both TRPA1 and TRPV regulate anandamide-induced scrunching in *D. japonica*. It remains to be determined which other receptor(s) may be responsible for the other scrunching inducers, including capsaicin, low pH and noxious heat. Importantly, because scrunching in response to capsaicin, heat or pH were not affected by knockdown of either *DjTRPA* or *DjTRPVab*, these genes are likely not responsible for the general scrunching response but rather mediate sensing of specific stimuli.

## Conclusions

Combining the results presented here with our previous studies of scrunching allowed us to partially decipher the molecular mechanisms responsible for sensing noxious stimuli in planarians. In this and our previous work, we found that TRPA1 and TRPV channels, as well as the big potassium channel SLO-1 (62), are involved in inducing scrunching in response to specific stimuli. Using RNAi, we found that some inducers are specific to one of these pathways, such as AITC and H_2_O_2_ to TRPA1, while others, such as anandamide and amputation respectively, rely on potential overlapping functions of both channels or on other unidentified channels. Lastly, the mechanisms of some scrunching inducers, including capsaicin, low pH, and noxious heat remain elusive. Further complicating matters, we found species differences with several of the scrunching inducers. For example, while anandamide induced scrunching mediated by both DjTRPAa and DjTRPV in *D. japonica*, it induced peristalsis in *S. mediterranea*. These observations and the differences in sensitivity to H_2_O_2_ and HC-030031between the two species demonstrate that although these two species are closely related, caution must be used when extrapolating the pharmacological effects of one species to another.

It is striking that despite the existence of multiple induction routes, the dynamic features of scrunching are independent of the inducer, and hence of the sensing pathway. This suggests some form of signal integration occurring downstream of these receptors, as illustrated graphically in Fig 6, which represents our current understanding of the molecular mechanisms of scrunching induction.

**Fig 6.**
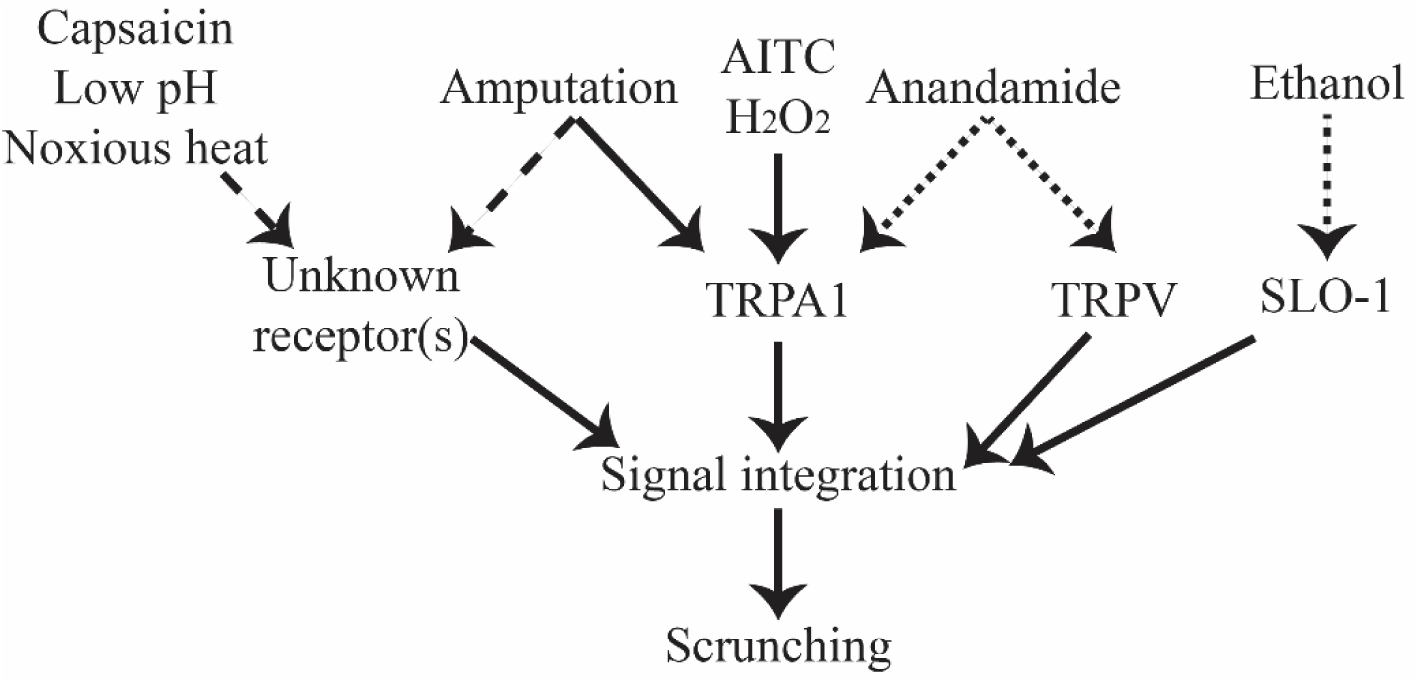
Overview of our current understanding of mediators of scrunching induction. Solid lines indicate that direct connections have been experimentally shown. Dotted lines indicate inducers which were only found to induce scrunching in one of the two species. Dashed lines are hypothesized connections.

Signal integration could occur at the neuronal level, raising the question of which parts of the planarian nervous system are involved. Our previous results (4) have shown that tail pieces which lack a brain are still capable of scrunching in response to some stimuli, albeit much more rarely. This would suggest that the brain, while not required for scrunching, plays an important role in achieving consistent induction.

Our pharmacological studies revealed that several agonist-antagonist pairs (AITC/HC-030031 and capsaicin/SB-366791) either both triggered scrunching and/or were unable to pharmacologically rescue the scrunching phenotype. These results were surprising given that in other systems, including invertebrates such as the medicinal leech and schistosomes, the antagonists work as expected to inhibit the action of the agonists (16,39,57). In contrast, in the two planarian species studied here, we found that both a TRPA1 agonist and antagonist induced scrunching, with potentiating effects when co-exposed in both planarian species. Similarly, although the TRPV antagonist SB-366791 did not induce scrunching alone, it also potentiated capsaicin-induced scrunching in *D. japonica*. Only RNAi against the target genes allowed us to suppress scrunching in response to specific chemical inducers. One possible explanation for these findings is that the planarian sensory system is highly sensitive to any deviation from normal and that scrunching is a default downstream response to system perturbations. However, the observed species differences demonstrate that scrunching is not always triggered, in agreement with our previous findings that ethanol, but not methanol, trigger scrunching (62). This argues that scrunching is a specific response, whose regulation, despite the progress made in this work, remains poorly understood.

Understanding the molecular mechanisms controlling the scrunching gait, from initiation to execution will require systematic studies of these different aspects using chemical and/or molecular approaches as presented here. The observed complexity and myriad of pathways involved in scrunching initiation reported here may explain why scrunching is a sensitive readout of neurotoxicity (8) and gaining a deeper understanding of its regulation will allow for more mechanistic studies of potential toxicants in the future using the planarian system. Moreover, by understanding the extent that aspects of nociception are conserved (or not) across species will provide better informative context to understand species-specific sensitivity differences and provide insight into the important mechanisms regulating noxious stimuli and pain sensation.

## Acknowledgments

The authors thank Cambria Neal, Arya Dadhania, Kelson Kaj, Angel Leu, Addam Debebe, Eileen Tsai, and Yingtian He for help with planarian care and experiments, Dr. Marco Gallio for the TRPA1 plasmid and discussions, Oscar Arenas for discussions of the AITC experiments, and Dr. William Kristan for discussions and comments on the manuscript.

