## Supplemental Information for "Pharmacological or genetic targeting of Transient Receptor Potential (TRP) channels can disrupt the planarian escape response"

### Supporting Information

**S1 Table. Maximum elongation and speed of scrunching are dependent on AITC concentration.** Scrunching parameters for *D. japonica* and *S. mediterranea* exposed to 50, 75, or 100 µM AITC, denoted as mean ± standard deviation. For each concentration, planarians were observed to scrunch with the parameters listed within the first minute in the bath. * denotes p < 0.05 and ** denotes p < 0.01 significance level compared to 50 µM AITC given by a two-tailed t-test. ^ denotes p < 0.05 and ^^ denotes p < 0.01 significance level compared to amputation given by a two-tailed t-test. [a]Amputation data are previously published values (4), provided for reference.

| **Species** | **Induction** | Frequency (cycles $s^{-1}$) | Maximum elongation | Speed (body length $s^{-1}$) | **N=** |
|---|---|---|---|---|---|
| *D. japonica* | 50 µM AITC | 0.72 ± 0.08 | 0.52 ± 0.04 | 0.37 ± 0.04 | 9 |
| *D. japonica* | 75 µM AITC | 0.78 ± 0.13 | 0.56 ± 0.06 | 0.44 ± 0.09* | 9 |
| *D. japonica* | 100 µM AITC | 0.92 ± 0.06**, ^ | 0.60 ± 0.03**,^^ | 0.55 ± 0.05**, ^^ | 8 |
| *D. japonica* | Amputation[a] | 0.70 ± 0.27 | 0.50 ± 0.08 | 0.34 ± 0.12 | 15 |
| *S. mediterranea* | 50 µM AITC | 0.33 ± 0.02 | 0.43 ± 0.05 | 0.14 ± 0.01 | 5 |
| *S. mediterranea* | 75 µM AITC | 0.42 ± 0.14 | 0.49 ± 0.02* | 0.21 ± 0.07* | 5 |
| *S. mediterranea* | 100 µM AITC | 0.41 ± 0.09 | 0.51 ± 0.12* | 0.22 ± 0.02** | 5 |
| *S. mediterranea* | Amputation[a] | 0.40 ± 0.09 | 0.44 ± 0.09 | 0.17 ± 0.09 | 77 |

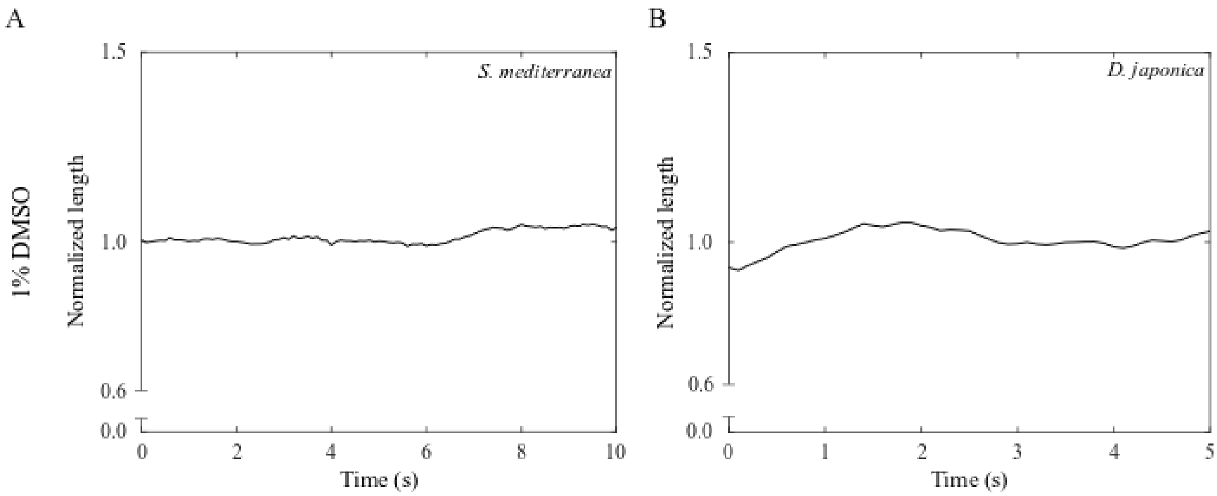


**S1 Fig. 1% DMSO does not induce scrunching in either *S. mediterranea* or *D. japonica.*** Representative length versus time plot for wildtype (A) *S. mediterranea* or (B) *D. japonica* planarians in 1% dimethyl sulfoxide (DMSO) (N=10). Planarians were exposed to 1% DMSO by directly pipetting 100 μL.

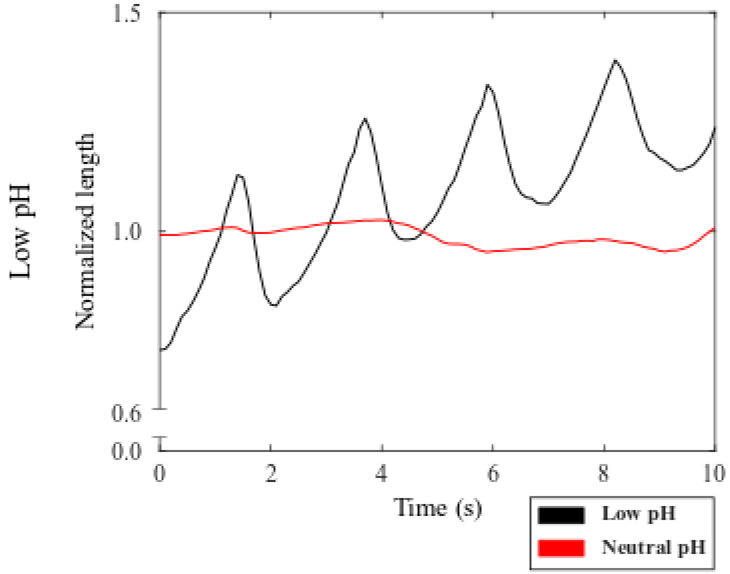


**S2 Fig. *S. mediterranea* scrunch in response to low pH exposure.** Representative length versus time plot for wildtype *S. mediterranea* planarians in Instant Ocean water at neutral pH (red, N=5) and pH 2.7 (black, N=10). Scrunching was induced by directly pipetting 100 µL onto planarians

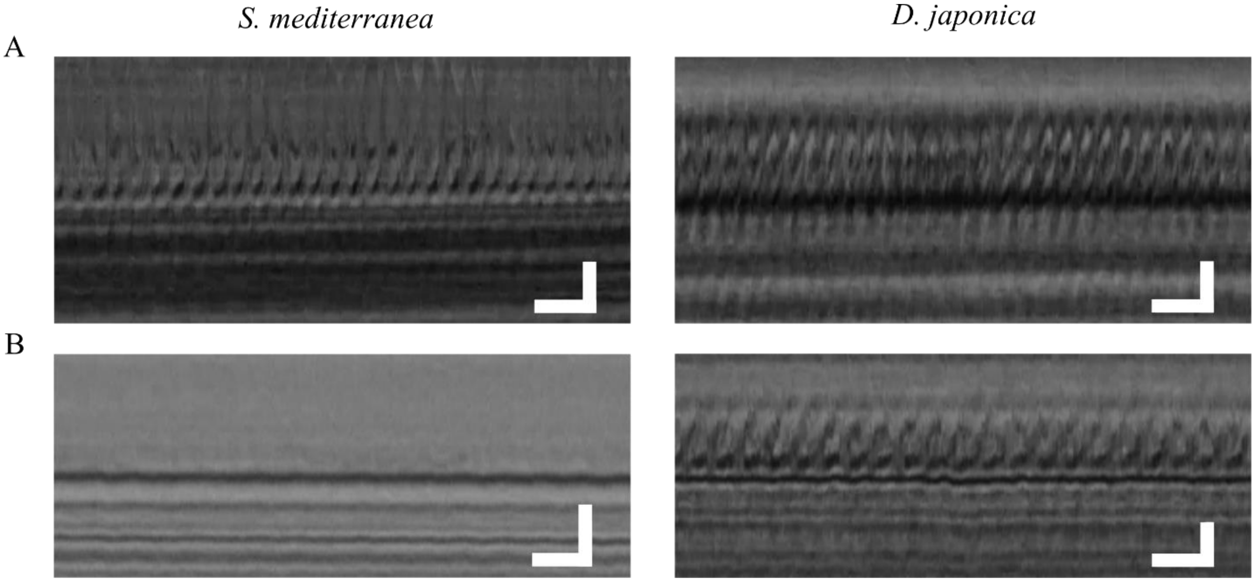


**S3 Fig. Anandamide impairs cilia beating in *S. mediterranea* but not *D. japonica*.** Representative (N=3/3) 1 s kymograph of cilia beating for *S. mediterranea* (left) and *D. japonica* (right). (A) Controls in planarian water and (B) when exposed to 100 μM anandamide for 5 minutes. Notice that cilia beating is almost completely lost in *S. mediterranea* while cilia beat normally in *D. japonica*. Scale bar shows 0.1 s horizontally and 1 μm vertically.

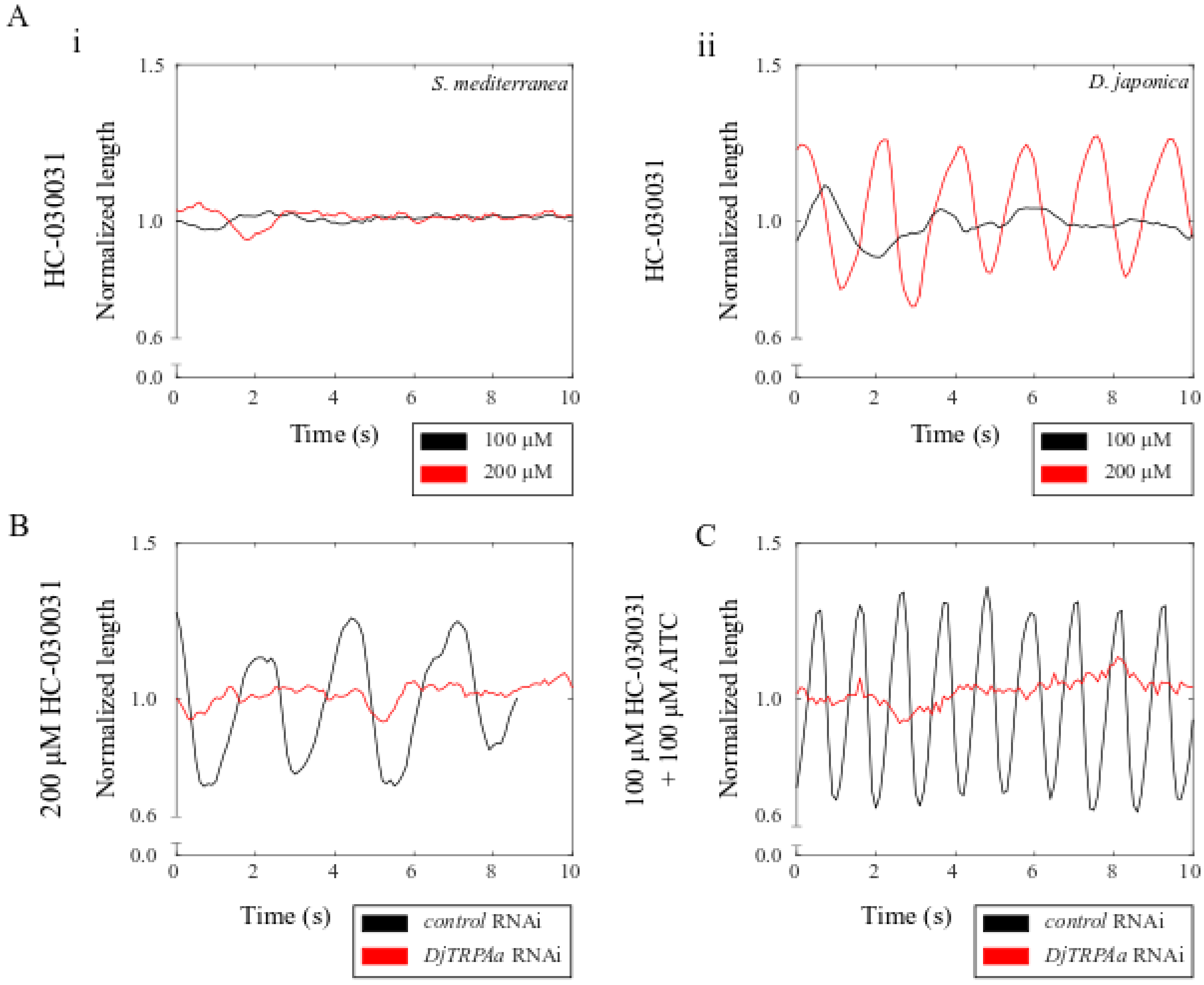


**S4 Fig. 200 µM HC-030031 induction of scrunching in *D. japonica* is dependent on TRPA1.** (A) Representative length versus time plot for wildtype (i) *S. mediterranea* and (ii) *D. japonica* planarians in 100 µM (black) and 200 µM (red) HC-030031. Plots are representative of N=10. (B) Representative length versus time plot for *D. japonica control* RNAi (N=13) and *DjTRPAa* RNAi (N=11) planarians in 200 µM HC-030031. (C) Representative length versus time plot for *D. japonica control* RNAi (N=8) and *DjTRPAa* RNAi (N=9) planarians in 100 µM HC-030031 + 100 µM AITC.

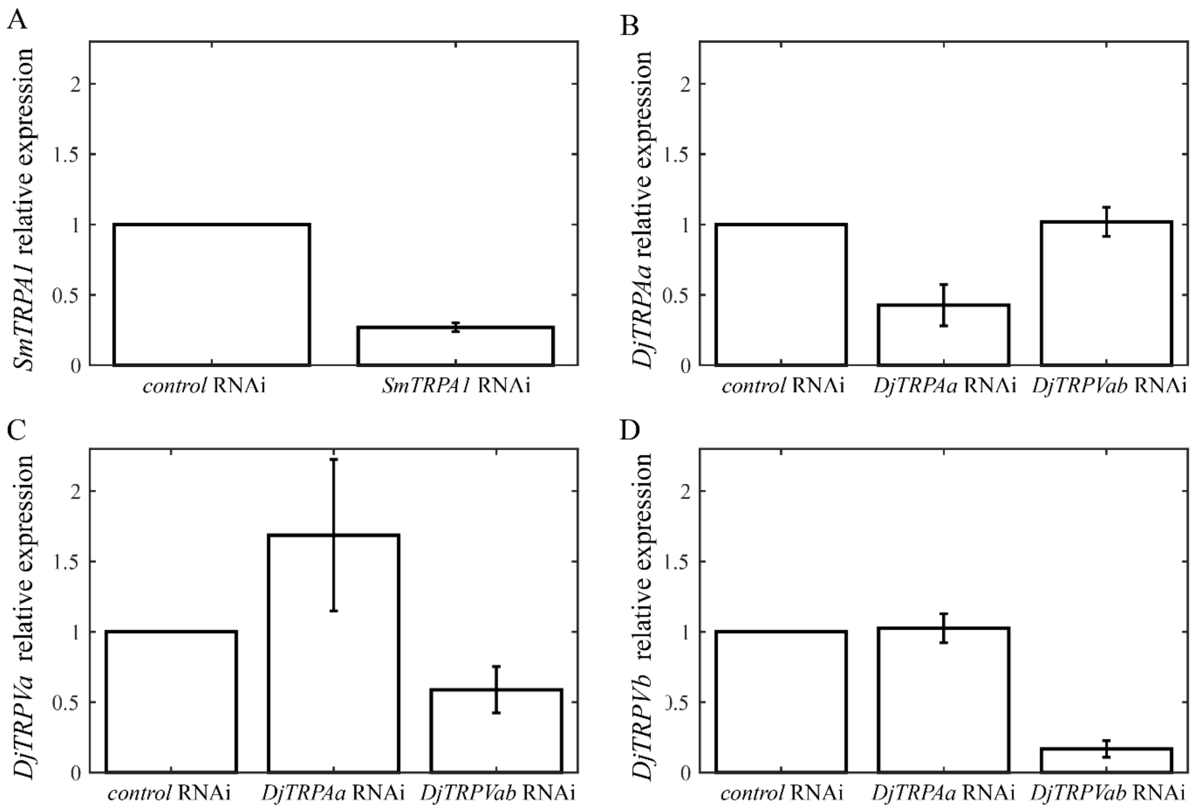


**S5 Fig. Confirmation of RNAi knockdown by qRT-PCR.** (A-D) Relative expression of (A) *SmTRPA1*, (B) *DjTRPAa*, (C) *DjTRPVa* and (D) *DjTRPVb* in the respective RNAi populations compared to the *control* RNAi population in that species. Data are shown as the mean of two biological replicates (each including 3 technical replicates). Error bars represent the standard error.

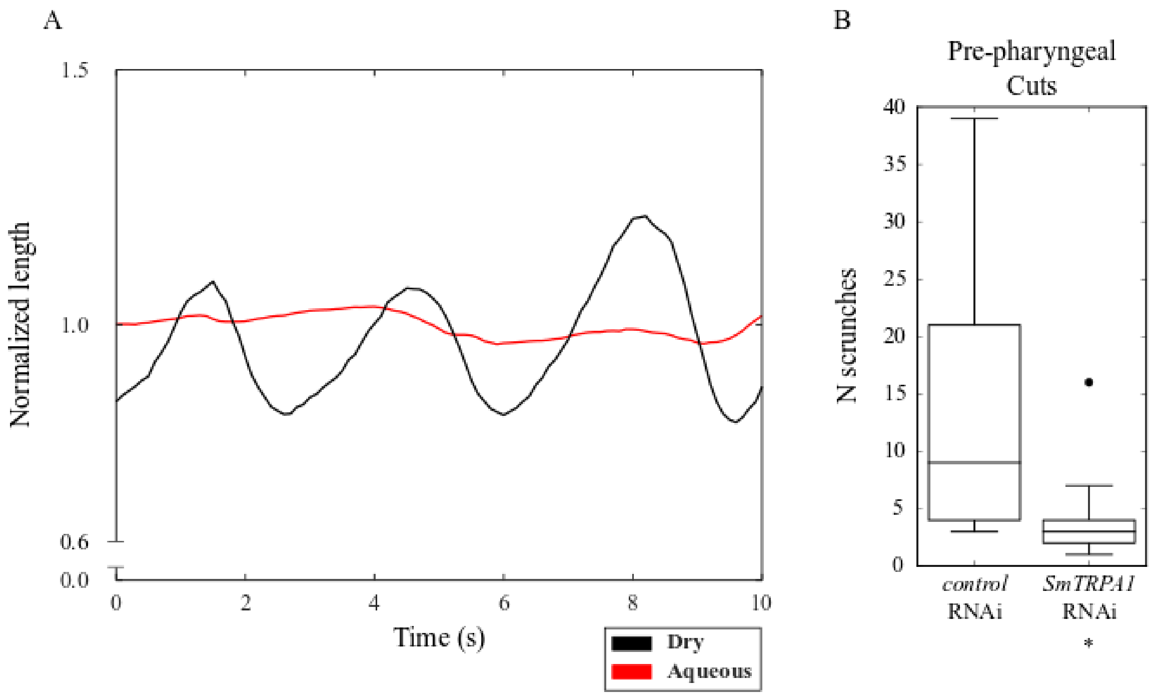


**S6 Fig. SmTRPA1 mediates scrunching in response to amputation.** (A) Representative length versus time plot for wildtype *S. mediterranea* planarians in an aqueous (red, N=5) and dry (black, N=10) environment, created by placing the planarians on wet filter paper as in (10). (B) Distribution showing median and quartiles of the number of scrunches directly following amputation in *control* RNAi (N=21) and *SmTRPA1* RNAi (N=24) planarians. * denotes $p < 0.01$ significance from *control* RNAi given by a two-tailed t-test.

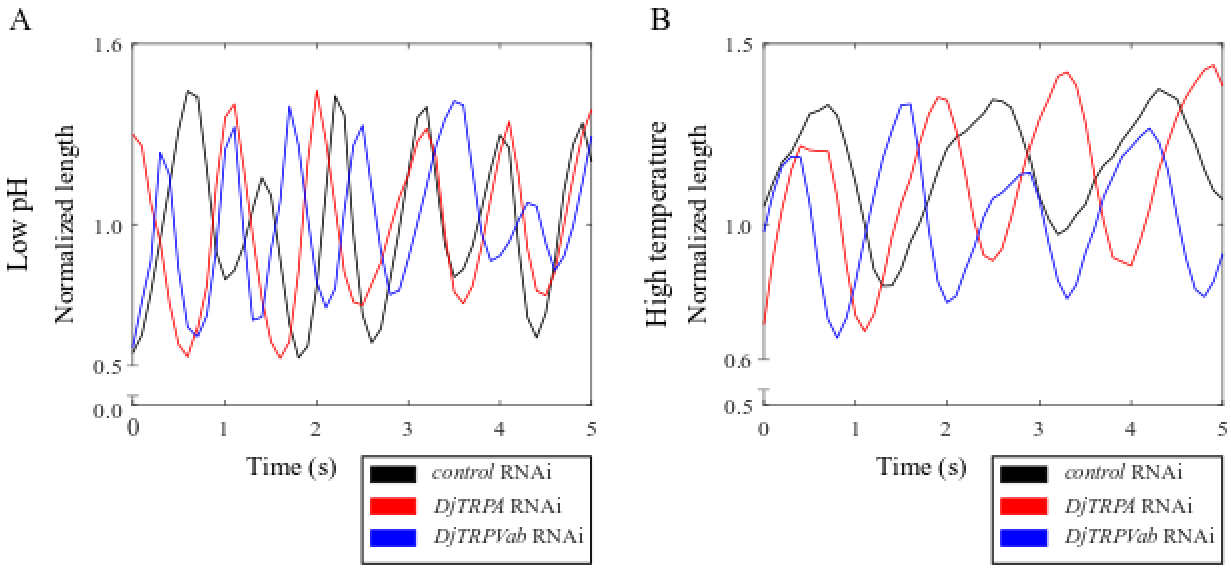


**S7 Fig. Heat and low pH sensing are not significantly impaired in *DjTRPAa* or *DjTRPVab* RNAi planarians.** (A-B) Representative oscillation plots for *control*, *DjTRPAa*, and *DjTRPVab* RNAi planarians exposed to (A) pH 2.7 and (B) 65°C IO water via pipette. No significant differences in scrunching induction are seen in any of the conditions. N=10 for all conditions.

**Supplementary Movies**

**S1 Movie. *D. japonica* and *S. mediterranea* planarians in 165 µM capsaicin.** Comparison of *D. japonica* and *S. mediterranea* behaviors in 165 µM capsaicin. Planarians exhibit vigorous head shaking and turning as well as scrunching. Movie is recorded and played at 10 fps.

**S2 Movie. *D. japonica* exposed to 50 µM AITC alone and with 100 µM HC-030031.** First 50 s of exposure of *D. japonica* to 50 µM AITC alone or co-administered with 100 µM HC-030031. Movie is recorded and played at 10 fps. Scale bar: 1 cm.

**S3 Movie. *S. mediterranea* exposed to 50 µM AITC alone and with 100 µM HC-030031.** First 50 s of exposure of S. *mediterranea* planarians to 50 µM AITC alone or co-administered with 100 µM HC-030031. Movie is recorded and played at 10 fps. Scale bar: 1 cm.

**S4 Movie. *D. japonica control* and *DjTRPAa* RNAi planarians in 100 µM AITC.** Behavior of *D. japonica control* RNAi and *DjTRPAa* RNAi in 100 µM AITC during the first 30 seconds of exposure. *Control* RNAi planarians scrunch whereas *DjTRPAa* RNAi planarians exhibit rapid head turning but lack a scrunching response. Movie is recorded and played at 10 fps. Scale bar: 1 cm.

**S5 Movie. *S. mediterranea control* and *SmTRPA1* RNAi *S. mediterranea* planarians in 100 µM AITC.** Behavior of *S. mediterranea control* RNAi and *SmTRPA1* RNAi planarians in 100 µM AITC during the first 30 s of exposure. Co*ntrol* RNAi planarians scrunch whereas *SmTRPA1* RNAi planarians glide. Movie is recorded and played at 10 fps. Scale bar: 1 cm.

**S6 Movie. *D. japonica control, DjTRPAa,* and *DjTRPVab* RNAi planarians in 165 µM capsaicin.** First 60 seconds of *D. japonica* RNAi populations when exposed to 165 µM capsaicin. Movie is recorded and played at 10 fps. Scale bar: 1 cm.

**S7 Movie. *D. japonica* behavior in 1 µM and 10 µM SB-366791.** Movie showing *D. japonica* behavior in SB-366791. *D. japonica* planarians exhibit no abnormal behaviors in 1 µM SB-366791 but display vigorous head turning at 10 µM SB-366791. Movie is recorded and played at 10 fps. Scale bar: 1 cm.

**S8 Movie. *D. japonica control, DjTRPAa,* and *DjTRPVab* RNAi planarians in 125 μM anandamide.** First 60 seconds of *D. japonica* RNAi populations when exposed to 125 μM anandamide. *Control* RNAi *D. japonica* planarians exhibit either scrunching or a non-scrunching behavior. Conversely, both *DjTRPAa* and *DjTRPVab* RNAi planarians show a significant decrease in both reactions. Movie is recorded and played at 10 fps. Scale bar: 1 cm.
